# EEG-based Decoding of Auditory Attention to Conversations with Turn-taking Speakers

**DOI:** 10.1101/2025.06.20.660726

**Authors:** Iris Van de Ryck, Nicolas Heintz, Iustina Rotaru, Simon Geirnaert, Alexander Bertrand, Tom Francart

## Abstract

**Objectives:** Auditory attention decoding (AAD) refers to the process of identifying which sound source a listener is attending to, based on neural recordings, such as electroencephalography (EEG). Most AAD studies use a competing speaker paradigm where two continuously active speech signals are simultaneously presented, in which the participant is instructed to attend to one speaker and ignore the other speaker. However, such a competing two-speaker scenario is uncommon in real life, as speakers typically take turns rather than speaking simultaneously. In this paper, we argue that decoding attention to conversations (rather than individual speakers) is a more relevant paradigm for testing AAD algorithms. In such a conversation-tracking paradigm, the AAD algorithm focusses on switching between entire conversations, resulting in less frequent attention shifts (ignoring turn-taking within conversations), thereby allowing for more relaxed constraints on the decision time.

**Design:** To test AAD performance in such a conversation-tracking paradigm, we simulated a challenging restaurant scenario with three simultaneous two-speaker conversations, which were podcasts presented in front of the listener and in the back left and back right of the room. We conducted an EEG experiment on 20 normal-hearing participants to compare the performance of AAD in the commonly used competing speaker paradigm with two speakers versus the conversation tracking paradigm with 2 or 3 conversations, each containing two turn-taking speakers.

**Results:** We found that AAD, using stimulus decoding, worked well under all experimental conditions, and that the accuracy was not influenced by the direction of attention, the proximity to the target conversation, or the presence of within-trial attention switches (versus a condition with sustained attention). Given the challenging scenario, we probed for the participants’ listening experience and found a correlation between the neural decoding performance and the perceived listening effort and self-reported speech intelligibility. To gain insight into the speech intelligibility of the participants in our setup, they performed a speech-in-noise test (Flemish matrix sentence test), but we did not find a correlation between the speech intelligibility performance and the AAD performance.

## 1. Introduction

Auditory Attention Decoding (AAD) is a technique used to determine which sound source a listener is paying attention to by analysing neural signals, which are typically measured using brain imaging techniques such as electroencephalography (EEG) or magnetoencephalography (MEG) (Biesmans et al., 2017; Ding & Simon, 2012; Mesgarani & Chang, 2012; J. A. O’Sullivan et al., 2015). When we pay attention to a specific auditory signal, such as a person speaking in a noisy environment, the neural activity phase-locks with certain features of that sound, such as its fundamental frequency (F0) or to the slowly fluctuating speech envelopes (Coffey et al., 2019; Howard & Poeppel, 2010; Luo & Poeppel, 2009; Pasley et al., 2012; Peelle et al., 2013). This synchronization between brain activity and sound is higher for an attended (speaker) than for an unattended speaker, which is used by AAD systems to decode selective attention (Biesmans et al., 2017; J. A. O’Sullivan et al., 2015). Many algorithms have been developed for AAD and are being developed and evaluated further (Biesmans et al., 2017; Das et al., 2016, 2017; Geirnaert et al., 2019, 2020, 2021, 2022; Han et al., 2019; Heintz et al., 2025; J. A. O’Sullivan et al., 2015; Rotaru et al., 2024; Shi et al., 2025; Tanveer et al., 2024; Van Eyndhoven et al., 2017; Wang et al., 2021; Wong et al., 2018; X. Xu et al., 2024; Z. Xu, Bai, Zhao, Hu, et al., 2022; Z. Xu, Bai, Zhao, Zheng, et al., 2022). Furthermore, this technology has potential applications in neuro-steered hearing aids, e.g., enabling devices to amplify only the attended sound (Das et al., 2020; Van Eyndhoven et al., 2017), as well as other applications in domains such as art, health, education, games, neuromarketing, etc. (Belo et al., 2021).

To apply AAD in practical applications, such as neuro-steered hearing aids (Van Eyndhoven et al., 2017), the AAD decoder must be able to work well in complex, realistic listening scenarios. Most AAD studies use a paradigm with two continuously active competing speakers, in which the participant has to attend to one speaker and ignore the other speaker (Biesmans et al., 2017; Das et al., 2016, 2017, 2018; de Cheveigné et al., 2018; Holtze et al., 2021; J. A. O’Sullivan et al., 2015; Wang et al., 2021). However, this setting with sustained attention to one of two competing speakers is not very realistic or common in everyday life, since speakers usually engage in conversations, in which they take turns. The listener would then frequently and rapidly switch attention between these turn-taking speakers, which is hard to track with state-of-the-art AAD algorithms.

Therefore, we question the traditional paradigm with two competing speakers as a validation setting for AAD. To make AAD practical for everyday applications, it is essential to evaluate its performance in more challenging and realistic environments with multiple conversations involving turn-taking speakers. We argue that - in practice - it is sufficient to track attention to a conversation rather than to individual speakers, such that tracking rapid and frequent attention switches within a conversation is not necessary. By treating the speech of both speakers within a conversation as a single entity, the algorithm focuses on switching attention between entire conversations rather than individual speakers within the conversation. This reduces the frequency of attention shifts and allows for more relaxed constraints on the AAD decision window length, i.e., the amount of EEG data needed for the AAD algorithm to make a sufficiently accurate decision on the current attention state.

Up to our knowledge, the only study involving a conversation-tracking paradigm for AAD is the one in Choudhari et al. (2024), where AAD was performed using intracranial (invasive) EEG (iEEG) recordings. They recorded iEEG in three neurosurgical patients, while they were asked to focus on one of two concurrent conversations with multiple talkers taking turns and continuously moving in space. Their system combined brain decoding with a binaural speech separation model that separated the speech of moving talkers while retaining their spatial locations. In addition, 24 (self-reported) normal-hearing participants participated in an online experiment in which they had to subjectively rate the proposed system. Those 24 subjects only participated in the audio perception experiment and did not participate in the EEG experiment. There was one condition with the system off, one condition with the system on where the separated speech was used for enhancement, and one condition with the system on where the original clean speech was used for enhancement. They found that their proposed system improved auditory perception, which included improved accuracy in detecting repeated words in the attended conversation, a decrease in the detection of repeated words in the unattended conversation, and a comparable localization across conditions. They also found that the proposed system improved auditory attention decoding and speech intelligibility, as well as reduced listening effort (Choudhari et al., 2024). Although the results were positive, some critical nuances should be considered. The AAD results are based on data from only 3 patients, which may not be representative for a larger population. Furthermore, the data is recorded with iEEG on neurosurgical patients. Generalisability to other more diverse groups, or to non-invasive neurorecording modalities such as EEG, can be limited. Furthermore, the experimental setup does not contain any intra-trial attention switches between the two conversations and may not completely reflect the complexity of everyday situations.

This paper advocates for a paradigm shift for evaluating AAD algorithms, from tracking individual speakers to decoding attention to conversations. In a conversation-tracking paradigm, the AAD algorithm focusses on switching between entire conversations rather than individual speakers. This approach naturally reduces the frequency of attention shifts (ignoring turn-taking within conversations) and allows for more relaxed constraints on the decision time, which is a more meaningful framework for evaluating AAD algorithms.

To validate this proposed paradigm, we performed an EEG experiment with a conversation-tracking paradigm in a realistic listening scenario. We designed four conditions (see Table 2) to answer the following questions. Is AAD as effective at decoding conversations as it is with single speakers? Is AAD capable of accurately decoding auditory attention in complex environments with three concurrent conversations?

## 2. Methods

### 2.1 Participants

This study includes 20 normal-hearing (≤ 25dB HL), young subjects (18 years and 10 months - 29 years and 4 months, with a median age of 20 years and 5 months). Eighteen subjects were female and two were male. All participants signed an informed consent at the time of arrival. The study was approved by the UZ Leuven/Research Medical Ethics Committee (KU Leuven) with reference S57102. At the beginning of the study, participants completed a questionnaire to verify inclusion and exclusion criteria, including age and medical history. Otoscopy was performed to check for visible abnormalities in the ear canal and the tympanic membrane. Hearing thresholds were assessed using pure tone audiometry, with a threshold of ≤ 25dB HL at all octave frequencies from 125 Hz to 8000 Hz. All participants were native Dutch (Flemish) speakers. Individuals with neurological disorders affecting cognition or attention, such as autism spectrum disorder (ASS), attention deficit (hyperactivity) disorder (AD(H)D), dyslexia, dyscalculia, or epilepsy, were excluded from the study.

### 2.2 Speech-in-noise test

To investigate a possible link between speech intelligibility and AAD performance, and to gain insight into the speech intelligibility of the participants in our setup, they performed a speech-in-noise test, the Flemish matrix sentence test (Luts et al., 2014). The sentences had a fixed grammatical structure, with a subject, verb, number, colour, and lastly an object. The participants had the choice between a fixed set of 10 words per category that were displayed on the screen in front of them. After hearing the sentence, participants had to click on the words they perceived or click ‘pass’ if they could not perceive any of the words.

The test was done in free field instead of headphones, to mimic our protocol setup in which the stimuli were also presented in free field (see subsection 2.3). We used the adaptive procedure, in which the sound level of the speech was constant (60 dBSPL) and the noise (standardized stationary speech-weighted) level was varied, based on the method of Wagener et al. (Wagener, Brand, et al., 1999a, 1999b; Wagener, Kühnel, et al., 1999) also known as the Oldenburg method, to get the speech-reception threshold (SRT). The SRT is the SNR at which 50% speech intelligibility is obtained. Speech and noise were spatially separated. The speech was presented from a loudspeaker in front of the participant, and the noise was presented from a loudspeaker above them. Presenting the noise from above the participant accounted for head-shadow effects and ensured consistent masking across frequencies. The matrix sentence test was standardised to administer the test with headphones or in free field without spatial separation. Therefore, we could not directly compare the results with the standardised norms, but our results can provide insights into speech intelligibility for our setup.

### 2.3 Apparatus

The participants were seated in a comfortable chair in front of 3 screens in a soundproof booth (see Figure 1).

**Figure 1.**
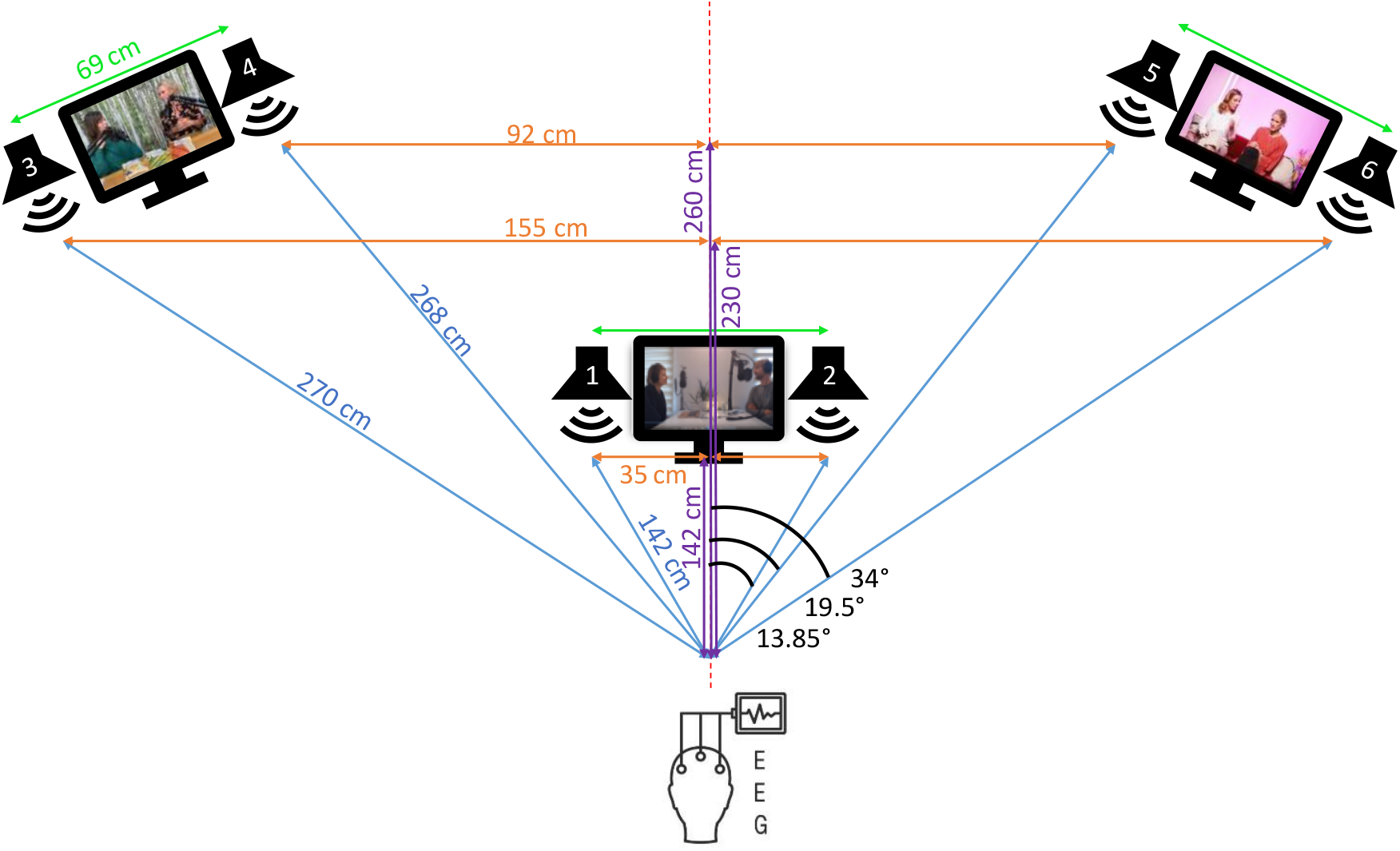
Visual representation of the setup. It also shows the positioning of the screens and loudspeakers (distance and angles) in the experimental setup in relation to the participant.

One screen was placed at the back left, one at the back right, and one right in the middle, in front of the participant. Next to each screen there were 2 loudspeakers. See Figure 1 for the exact position of the loudspeakers. Stimuli were presented using a Behringer (U-PHORIA UMC1820 audio-interface) sound card.

Participants were instructed to attend one of the speakers or conversations and ignore the other speakers. The attention cue was shown with a coloured rectangle around the videos (green = attended, red = unattended). Throughout the conditions, brain activity was recorded with 64-channel Biosemi ActiveTwo EEG. Participants were instructed to move as little as possible to avoid muscle artefacts in the EEG signals. Communication with the participants could be maintained via a microphone in the booth.

### 2.4 Stimuli

Stimuli were presented at 60dB A from the frontal loudspeakers and at 54.5 dB A from the speakers in the back. The intensity was calculated based on the distance from the participant to the loudspeaker to imitate the natural attenuation with distance. The sound level was calibrated at the position of the listener.

The stimuli of the condition with two competing speakers were randomly selected podcasts from the collection of ‘University of Flanders’ (Universiteit van Vlaanderen). The stimuli with conversations were randomly selected podcasts (with video) about various topics, each containing two turn-taking Flemish speakers without a strong dialect. When selecting the podcasts, we made sure that both speakers were always visible during the 10-minute video fragment. The videos were pre-processed with the program VSDC, to make sure they all had the same sampling frequency (44100 Hz), the intros were cut, and the videos were shortened to 10 minutes. We manually separated the voices of the podcast speakers into individual mono files using Audacity, allowing us to play each speaker through a different loudspeaker to mimic natural spatial separation between speakers in a conversation. The stimuli were further processed in Python^1^.

### 2.5 Conditions

The experiment contained 4 conditions and 10 minutes of data per trial. The different trials within each condition indicated which speaker or conversation was being attended to (i.e. the locus of attention). In the **2 Competing Speakers** condition, there were two simultaneous competing speakers and in the **2 Conversations** condition, there were two simultaneous conversations (four speakers turn taking in pairs) presented. The other two conditions involved three conversations. In the **Sustained Attention** condition, the participant attended to one of the three conversations throughout the condition. Lastly, in the **Attention Switch** condition, participants attended to one of the three conversations, and after 5 minutes they switched their attention to one of the other conversations. A visual representation of the different conditions is shown in Figure 2, with green indicating the attended speaker(s) and red the ignored/unattended speaker(s).

**Figure 2.**
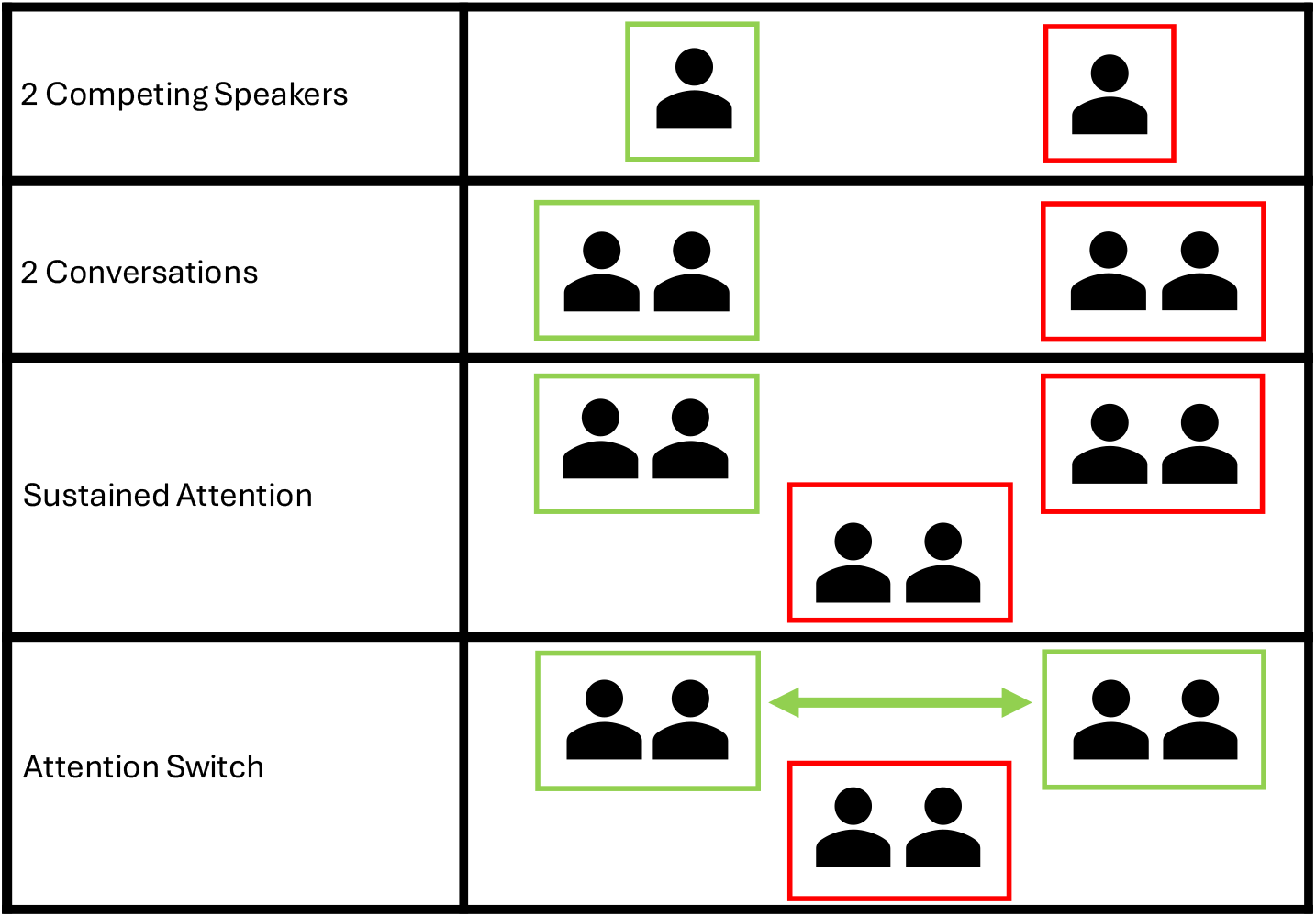
Visual representation of the different condition types. The green rectangle indicates the attended speakers/conversations, and the red rectangle represents the unattended speakers.

The order of the conditions and stimuli was randomised differently for each participant. The 3 trials of sustained attention and attention switch conditions always stayed together in one block, but the individual trials were randomized in order. See the overview of the experimental conditions in Table 1 and a visual representation in Figure 2. In total, the test sessions took between 3 and 3.5 hours, which yielded 80 minutes (8 trials of 10 minutes) of EEG data consisting of the 4 experimental conditions. Breaks were obligatory after 3 consecutive trials. More breaks were taken if the participant indicated that they felt tired or needed a break.

**Table 1:** Overview of the conditions.

| Condition | Content | Number of trials | Attended speaker(s) |
| --- | --- | --- | --- |
| 2 Competing Speakers (Reference) | Classic 2 speakers AAD | 1 | Left or Right speaker |
| 2 Conversations | 2 conversations instead of 2 speakers | 1 | Left or Right conversation |
| Sustained Attention (3 conversations) | 3 conversations – attention only to one of the three conversations | 3 | Conversation at the Front<br>Conversation on the Left<br>Conversation on the Right |
| Attention Switch (3 conversations) | 3 conversations – switch attention from one conversation to another conversation | 3 | Switch between Front & Left conversation<br>Switch between Front & Right conversation<br>Switch between Left & Right conversation |

**Table 2:**
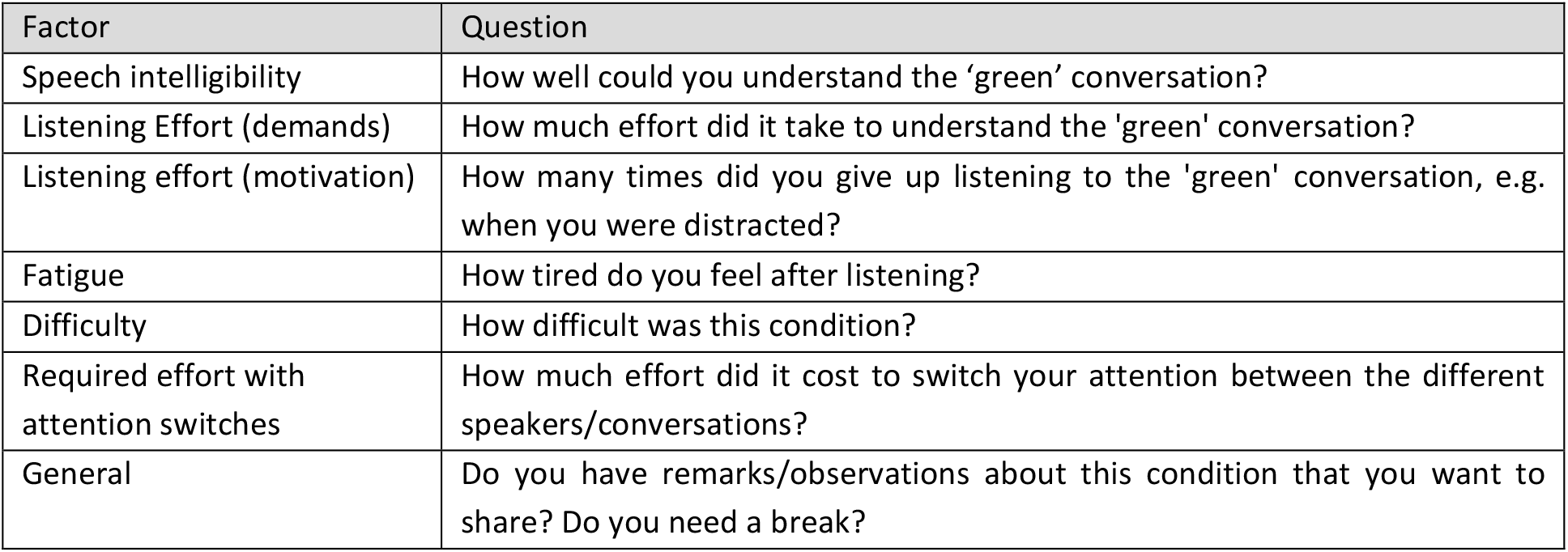
Behavioural questions assessing listening experience.

| Factor | Question |
| --- | --- |
| Speech intelligibility | How well could you understand the 'green' conversation? |
| Listening Effort (demands) | How much effort did it take to understand the 'green' conversation? |
| Listening effort (motivation) | How many times did you give up listening to the 'green' conversation, e.g. when you were distracted? |
| Fatigue | How tired do you feel after listening? |
| Difficulty | How difficult was this condition? |
| Required effort with attention switches | How much effort did it cost to switch your attention between the different speakers/conversations? |
| General | Do you have remarks/observations about this condition that you want to share? Do you need a break? |

### 2.6 Behavioural measures

After each trial, there were 2 multiple choice questions about the content of the story, to keep the participants’ attention and motivation high, followed by a few questions to assess the subjective experiences of the participant during the respective trial. Those questions evaluated speech intelligibility, listening effort, fatigue, difficulty, and effort in switching attention (see Table 2). These questions were displayed on the front screen, after which the participants answered verbally with a score out of 100 (%). In addition, there was a general question about their experience.

### 2.7 Signal processing

The EEG data was processed in MATLAB (R2023a) using the AAD algorithm proposed by Biesmans et al. (2017). To compare the performance between the four different conditions (eight trials), we report the accuracy of the AAD algorithm predicting the attended speaker.

In this study, we assume that the clean speech signals are available, and that we know which speech signals belong to the same conversation.

### Speech stimuli

The speech envelope was extracted from the original speech signal using the method described by Biesmans et al. (2017). This involved a gammatone filter bank to decompose the speech signals into sub-bands, followed by a power law compression applied to each sub-band. These sub-band envelopes are then summed to form a single envelope for auditory attention decoding (Biesmans et al., 2017). In the final step, the speech envelope was downsampled to 128 Hz.

### EEG processing

The EEG data were recorded at a sampling rate of 8192 Hz and processed per condition. Several preprocessing steps were applied to the EEG data:

1. Data were initially downsampled from 8192 Hz to 256 Hz using the MATLAB resample function with an anti-aliasing filter to reduce pre-processing time.
2. Eye-blink artefacts were estimated and removed using a Multichannel Wiener filter (MWF), based on the method outlined by Somers et al. (2018).
3. A bandpass Chebyshev filter was applied to retain frequencies between 1 and 9 Hz.
4. Following the bandpass filtering, all channels were re-referenced to the common average reference.
5. Finally, the data were downsampled again, to 128 Hz for further processing and to match the sampling rate of the speech envelope.

### AAD decoder

Similar to e.g. Han et al. (2019), J. O’Sullivan et al. (2017), Van Eyndhoven et al. (2017) we assume that AAD algorithms have access to clean, separated conversations. Thus, we make abstraction of the automated speech separation that would typically precede AAD.

For AAD, we used a backward least-squares linear decoder to reconstruct the speech envelope from the multichannel EEG (Biesmans et al., 2017). The reconstructed envelope is correlated with the original stimulus envelopes (Biesmans et al., 2017) and the one with the highest correlation is considered the attended one. A decision window length of 30 seconds was chosen, which refers to the duration of the window used to correlate the reconstructed envelope with the original stimulus envelopes.

There was a total amount of 80 minutes of data (8 trials of 10 minutes). A subject-specific decoder was trained using a **leave-one-trial-out (LOTO) Cross-Validation (CV)** method. For each iteration, the model was trained on data from 7 trials (70 minutes in total) and tested on the remaining trial (10 minutes). This process was repeated 8 times, so that each trial served once as the test set. The attended speaker was predicted for each segment, and prediction accuracies were then averaged per condition and per participant, which resulted in a single prediction accuracy (%) per condition per participant.

### 2.8 Statistical analysis

The statistical assumptions (normality, homogeneity) for the use of a parametric test were not met, so we chose non-parametric statistical tests. Statistical analysis was performed using R software (version 3.6.3). The significance level was set at α=0.05. The statistical methods chosen to investigate the effects on AAD decoding performance were the Wilcoxon Rank-sum test for a comparison between 2 groups and the Kruskal-Wallis test for more than 2 groups (Field et al., 2012).

To compare between all the conditions, given the differences in chance levels (50% for two speakers/conversations and 33.33% for three conversations), we also calculated an adjusted accuracy metric (similar to balanced or normalised accuracy; see (Brodersen et al., 2010)), and defined here as:

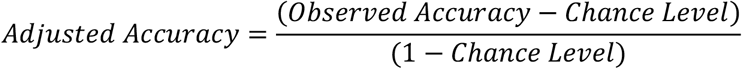

We note that all reported accuracies are the *unadjusted* ones, unless stated otherwise.

## Results

In the conversation study, we wanted to track attention to conversations instead of to individual speakers. Figure 3 shows the average AAD accuracy per condition for the 20 subjects.

**Figure 3.**
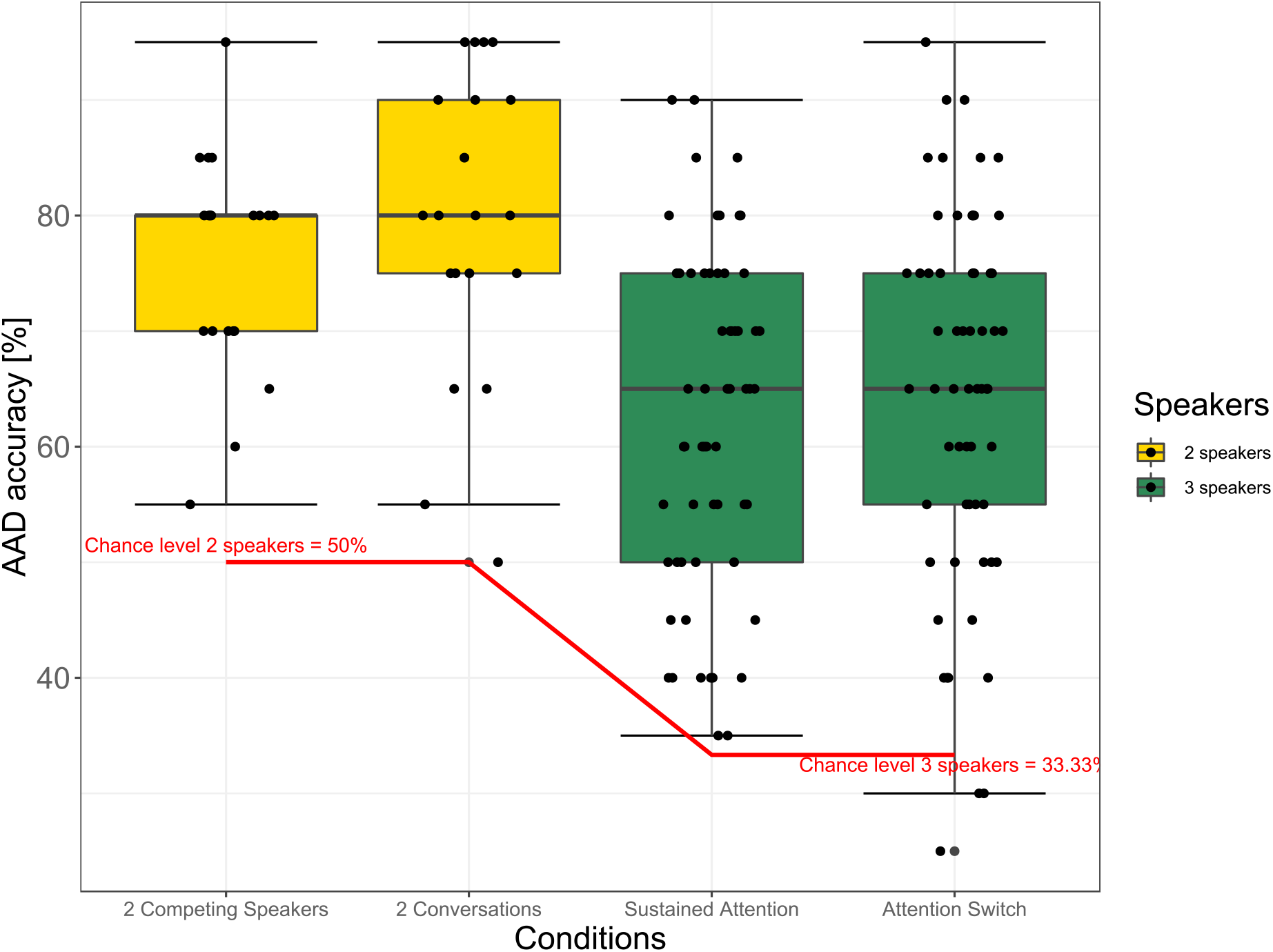
AAD results aggregated across all trials of each condition. Each dot represents a subject. The conditions with 2 competing speakers and 2 conversations are shown in yellow. The conditions with 3 speakers/conversations are shown in green (Sustained Attention and Attention Switch).

### General

In general, all conditions perform significantly better than chance (see Figure 3). For the conditions with 2 competing speakers and 2 conversations (with 2 speakers), this was significantly (< .0001) above 50% and for the Sustained attention and Attention switch with 3 speakers this was significantly (< .0001) above 33.33%. We calculated the significance per condition, taking into account their respective chance levels, on group level with the Wilcoxon Signed rank test (single sided).

### 2 conversations

The Wilcoxon Rank Sum (two sided) test demonstrated that there was no significant difference (p=0.296, 95% CI [-0.100 ; 0.050]) between decoding the auditory attention with two competing speakers and decoding two conversations.

### Switches

Wilcoxon’s rank sum test (two-sided) showed that there was no significant difference (p= 0.393, 95% CI [-0.050 ; 0.100] between decoding 3 conversations without and with intra-trial attention switches.

### Locus of attention

The Kruskal-Wallis rank sum test showed that there was no significant difference between the 3 trials within the Sustained Attention condition (p = 0.439, eff_size = -0.006, 95% CI [-0.03 ; 0.19]) and the Attention Switch condition (p = 0.511, eff_size = -0.012, 95% CI [-0.03 ; 0.16]), meaning that there was no significant difference in performance between decoding the attention from the frontal, right, or left speakers.

### 2 conversations versus 3 conversations

The mean adjusted accuracy (over subjects) per condition was 0.52 (SD 0.19) for the 2 Competing Speakers condition, 0.59 (SD 0.26) for the 2 Conversations condition, 0.44 (SD 0.22) for the Sustained Attention condition and 0.47 (SD 0.24) for the Attention Switch condition. The adjusted accuracies are presented in Figure 4. The interpretation of adjusted accuracies states that values between 0 (chance level) and 1 (perfect accuracy) represent proportional performance above chance. The adjusted values are surprisingly close across all conditions. This indicates that while raw performance drops in the 3 conversations conditions, relative to their difficulty, AAD still performs well. To test whether the adjusted accuracies differed significantly between conditions, we used a Kruskal-Wallis test. The results showed that there was no significant difference between the adjusted accuracies per condition (p = 0.092).

**Figure 4.**
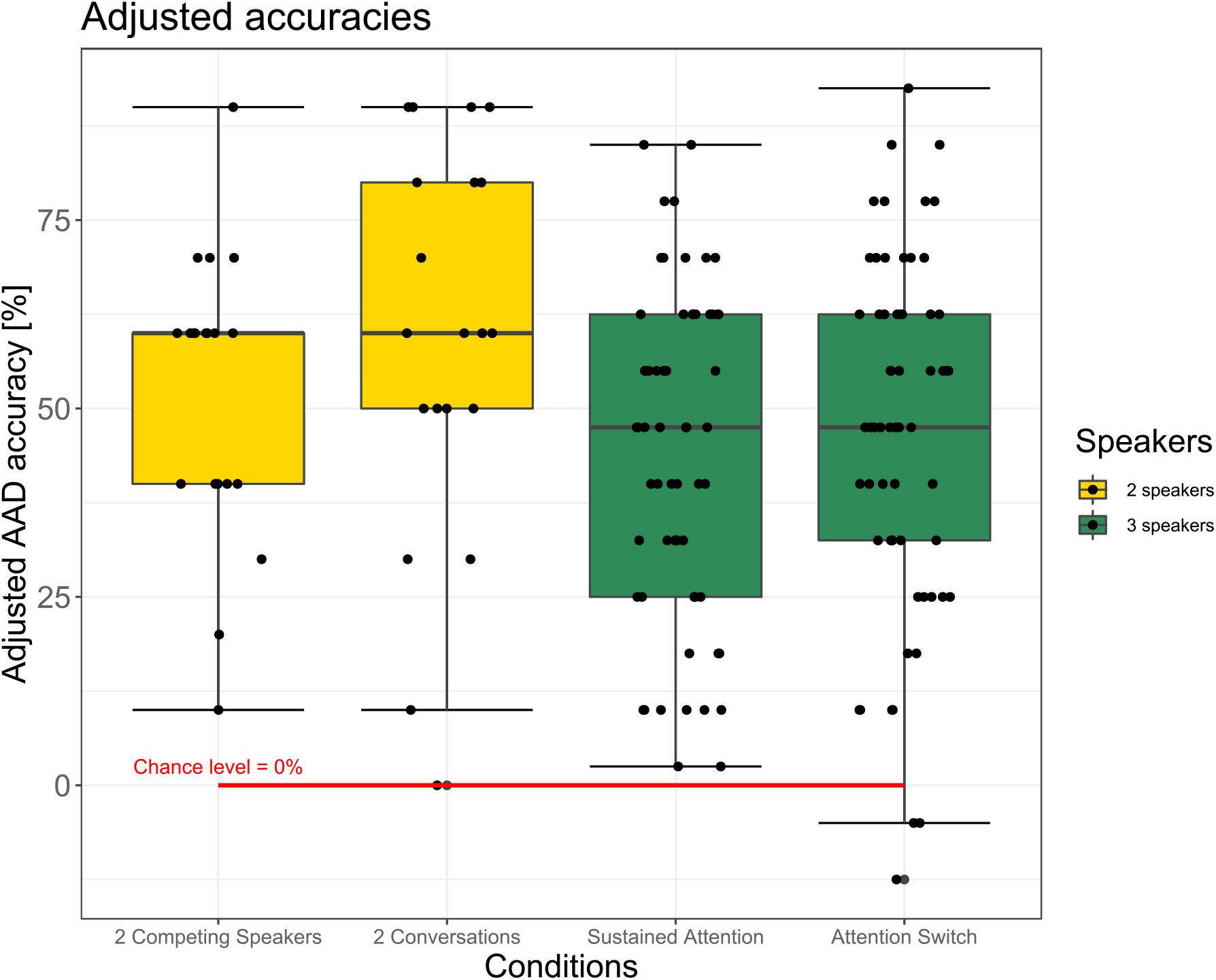
Adjusted accuracies per condition. AAD results aggregated across all trials of each condition. Each dot represents a subject. The conditions with 2 competing speakers and 2 conversations are shown in yellow. The conditions with 3 speakers/conversations are shown in green (Sustained Attention and Attention Switch).

### Behavioural measures

#### a. Behavioural questions

The participants answered behavioural questions about their experience after each trial. To examine the relationship between behavioural self-reports and neural AAD performance, the Spearman rank correlation test was used. We used the adjusted accuracies for this analysis because they take into account the different chance levels per condition. We tested over subjects and conditions to look at a general trend (not per condition or subject). The resulting p-values and Spearman’s rho coefficients are presented in Table 3. Scatterplots of the behavioural measures can be found in Supplementary Figure 1.

**Table 3:** Spearman rank correlation between behavioural questions and AAD performance (adjusted accuracies) (p-values and Spearman rho). An asterisk indicates significant correlations.

| Behavioural measures | p-value | Spearman's Rho |
| --- | --- | --- |
| **Speech intelligibility** | **0.005\*** | **0.256** |
| **Listening effort (demands)** | **0.017\*** | **-0.221** |
| Listening effort (motivation) | 0.068 | -0.169 |
| Fatigue | 0.7444 | 0.030 |
| Difficulty | 0.439 | -0.072 |
| Required effort with attention switches | 0.227 | -0.112 |

Among the behavioural measures, only self-reported speech intelligibility (p = 0.005 and ρ = 0.256) and listening effort (demands) (p = 0.017 and ρ = -0.221) showed a significant moderate correlation with (adjusted) AAD accuracy.

A covariance matrix (see Supplementary Figure 2) was computed to further explore potential interdependencies among the behavioural measures. A strong intercorrelation was observed between several behavioural measures (r up to 0.83), suggesting shared variance across constructs such as listening effort and difficulty.

#### b. Matrix sentence test (SRT)

The average SRT for our adapted test setup was -13.1 dB SNR across 20 normal-hearing participants, with an intersubject standard deviation of 2.33 dB. Intra-subject variability was 0.39 dB.

We used the Spearman rank correlation to calculate the correlations between the matrix scores per subject and their respective (unadjusted) AAD scores under the different conditions. No significant correlations were found for 2 Competing Speakers (p = 0.919, ρ = 0.024), 2 Conversations (p = 0.697, ρ = 0.093), Sustained Attention (p = 0.449, ρ = 0.100) and Attention Switch (p = 0.457, ρ = 0.098) conditions.

Additionally, we computed the correlation between the average (adjusted) AAD accuracy (per subject) across all conditions and the Matrix scores, with a spearman rank correlation and found p = 0.344, ρ = 0.223. Due to ties in the data (e.g. multiple identical accuracy scores), we also calculated the more robust Kendall’s Tau metric, which yielded a small positive correlation (τ = 0.168), which was also not statistically significant (p=0.319).

## Discussion

In this paper we introduced a conversation-based AAD validation paradigm with three simultaneous conversations, each consisting of two turn-taking speakers. This conversation-tracking paradigm emphasizes switching between entire conversations rather than individual turn-taking speakers, which results in less frequent and less abrupt attention shifts, thereby relaxing the speed requirement for AAD algorithms without compromising practical applicability. The results showed that all conditions scored significantly higher than chance. There was no significant difference between the conditions with 2 simultaneous speakers or with 2 conversations. This means that conversations can be decoded just like single speakers, even in this complex scenario. Also, once a third conversation is added to the setup/condition, the AAD performance is still significantly better than chance. We found that **3 competing conversations** can be decoded with an accuracy that is substantially better than chance. The adjusted accuracies were remarkably close across all conditions. This indicated that AAD performs equally well under different experimental conditions with different chance levels, even though raw performance drops in the 3 conversations conditions due to the higher number of classes.

AAD performance in our experimental conditions is similar to what we find in the literature for stimulus reconstruction with a linear decoder for a 2-speaker scenario with a window length of 30 seconds. For example Biesmans et al. (2017) found a detection accuracy of 81.5% for a window length of 30 seconds. In our study, AAD accuracies reached a mean of 76.0% (2 Competing Speakers) and 79.5% (2 Conversations). Schäfer et al. (2018) found that in 61.1% of the trials, their stimulus reconstruction algorithm was able to correctly classify the attention of the listener in a 4-speaker scenario. Note that the chance levels differed across studies, so direct comparisons should be interpreted with caution. For our conditions with sustained attention (3 conversations, each with 2 turn-taking speakers) the mean was 62.4% and for the attention switch condition it was 64.4%.

An effect of **switches** or **locus of attention** would be undesirable for practical applications of AAD, since we want AAD to perform well across all different kinds of real-life situations. Reassuringly, we found no significant differences between the conditions with and without a switch of attention. Similarly, no significant differences were observed between the 3 trials (front, right or left attended) within either the Sustained Attention or Attention Switch conditions. This is encouraging since in the trials we manipulated both the distance and the location of the speakers. Apart from switches or the locus of attention, other factors could influence AAD performance, which can be explored in future research. While some factors have already been partially explored (Das et al., 2016, 2018; Van de Ryck et al., 2025), many remain to be examined. These include both factors that are related to the speakers or to the listener, such as the content of the podcasts, participant’s motivation, levels of fatigue, educational background, cognition, age, degree of hearing loss, voice familiarity, topic familiarity, level of engagement (passive vs active), and listening effort (Van de Ryck et al., 2025).

To evaluate speech intelligibility in our set-up and compare it with AAD performance, we conducted a speech-in-noise test. The typical performance, that is, the average SRT of normal-hearing subjects, on the **matrix sentence test** is -9,5 dB SNR with an intersubject standard deviation of 0.8 dB (Luts et al., 2014). Using our adapted test protocol, we obtained an average SRT of -13.1 dB SNR across 20 normal-hearing participants, with a noticeably higher intersubject standard deviation of 2.33 dB. This shows a substantially larger spread between individuals, which is almost three times larger than the normative data. This suggests higher variability in speech intelligibility among participants in our test setup.

Despite this, intra-subject variability was 0.39 dB, which we consider acceptable and within expected limits for adaptive speech-in-noise testing.

Furthermore, we found no correlation between speech intelligibility performance on the Matrix test and AAD performance. Scores on the speech-in-noise task did not predict or relate to the brain scores.

### Behavioural measures

The participants answered 7 behavioural questions after each condition. Only self-reported speech intelligibility (p = 0.005 and ρ = 0.256) and listening effort (demands) (p = 0.017 and ρ = -0.221) showed a moderate significant negative correlation with (adjusted) AAD accuracy, suggesting that the more effort a participant reported, the lower their AAD accuracy. In addition, self-reported (subjective) speech intelligibility reflected the performance on objective measured speech intelligibility. The higher the speech intelligibility score the higher the AAD accuracy.

With a covariance matrix we further evaluated potential redundancies between the behavioural measures. If participants indicated that the listening experience under a condition was difficult, they often also indicated this in the questions regarding listening effort. Therefore, the questions might have measured a shared underlying construct such as mental load. This indicates potential redundancy or overlap between the different behavioural measures.

### Limitations of the study

Our study aimed to create a more realistic listening scenario to test the performance of a conversation-tracking AAD paradigm. We simulated a restaurant scenario with three simultaneous conversations. However, a more realistic simulated scenario might also include reverberation and background noise (Picou et al., 2019; Puglisi et al., 2021), and a larger number of conversations.

This study assumes access to clean speech signals and prior knowledge of their conversational groupings, which is a challenge that remains to be addressed in hearing aid systems.

Future research might explore other populations than our homogenous group of young normal-hearing subjects, including individuals of various ages and with different degrees of hearing loss. These diverse groups are particularly relevant as they represent the eventual users of neuro-steered aids.

## Conclusions

The majority of the AAD studies have focused on scenarios with two competing speakers in which participants were required to attend to one speaker while ignoring the other speaker. We questioned this classic approach and showed that it would be more relevant to switch to a conversation-tracking paradigm for future validation of AAD algorithms in realistic scenarios to improve the ecological validity of AAD research. Tracking entire conversations rather than individual speakers better matches natural social interactions. In addition, in real-life situations, more than two speakers frequently talk at the same time (cocktail party scenario).

We tested our conversations-tracking paradigm in a realistic, acoustically complex scenario with three simultaneous two-speaker conversations, using three screens and six loudspeakers in a soundproof test booth. We collected EEG data to investigate AAD performance under four experimental conditions. Twenty young normal hearing subjects were tested, and we found that an AAD algorithm, focused on tracking conversations rather than individual speakers, performed well across our experimental conditions within an acoustically complex scenario, which supported our claim. These are promising results for the future of AAD in practical applications.

## Author Statement

Tom Francart, Alexander Bertrand and I.V.d.R. designed the experiment (protocol and practical set-up). Iustina Rotaru helped with the pre-processing of the stimuli in Matlab and the Python scripts to run the experiment. Nicolas Heintz contributed to the analysis of the EEG data and adapted and applied the AAD decoders. All authors contributed to the discussion of the results and implications of the research. I.V.d.R. performed the experiments, performed the statistical analysis, and wrote the paper. All co-authors reviewed the paper.

## Acknowledgments

The authors would like to thank Liese Branders for helping with data acquisition, selection of the podcasts and composition of the multiple-choice questions. We would also like to thank all the subjects for their participation in the study.

## Declaration of generative AI and AI-assisted technologies in the writing process

During the preparation of this work the author used ChatGPT to rephrase sentences in academically correct English. After using this tool, the author reviewed and edited the content as needed and takes full responsibility for the content of the published article.

## Footnotes

1 The stimuli were each normalised to -27 dB RMS (by excluding the silences parts). Then, we determined the time points where the 2 voices from a podcast overlapped, to later apply a -3dB correction. This was done because the audio of each podcast was split into two channels corresponding to the two voices, left and right). When those voices overlapped, both voices were kept on both the left and right channel. Hence, when both channels were played together, the loudness was higher because the overlapped voices were played simultaneously from both channels. To avoid this amplification effect, a - 3dB correction for the voice overlapping sections was needed.

