## Supplementary figures and images for "EEG-based Decoding of Auditory Attention to Conversations with Turn-taking Speakers"

### Supplementary Figure 2

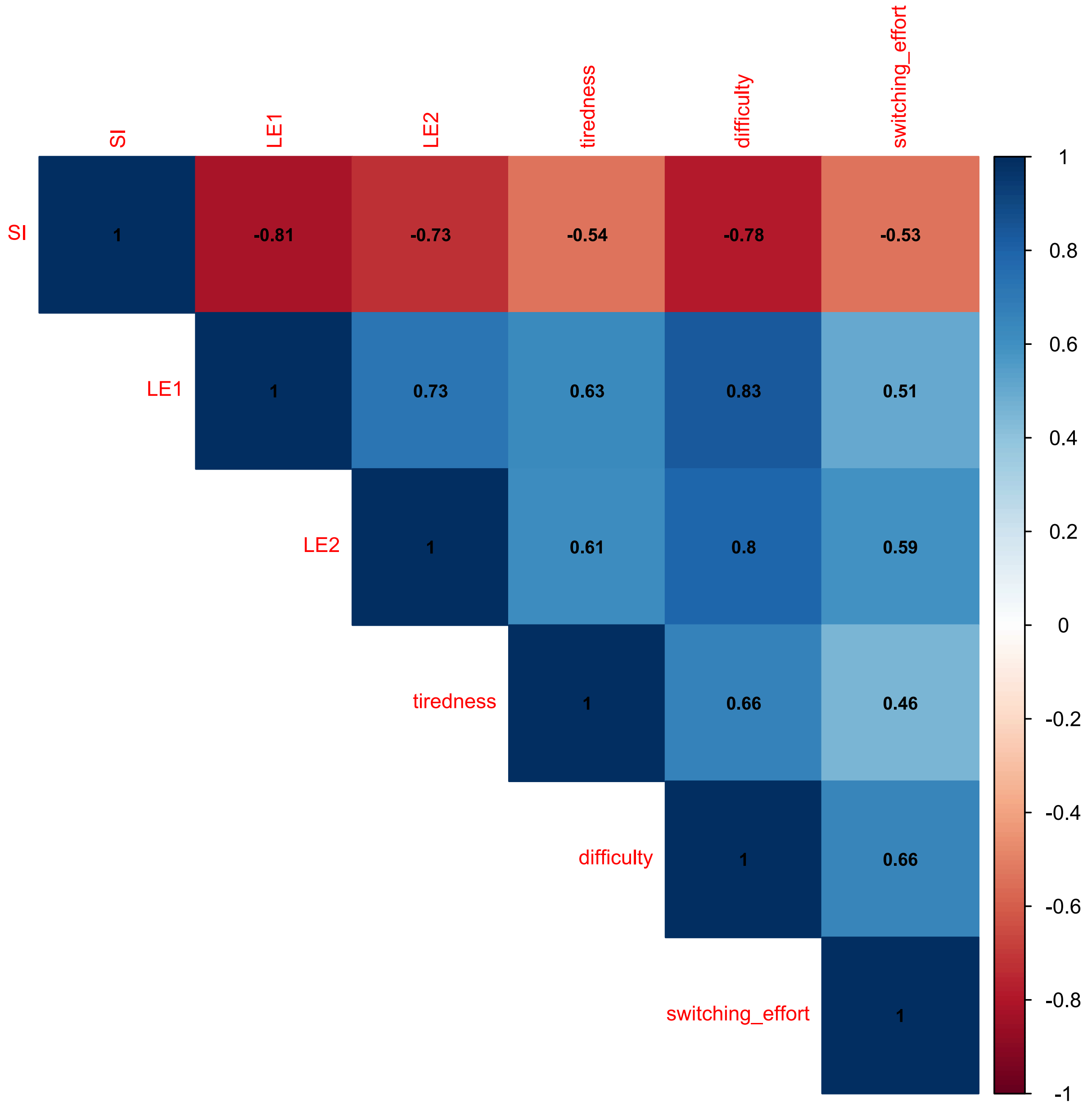


*Supplementary Figure 2: Covariance matrix of the behavioural measures*
