## Supplementary Figure 1 for "EEG-based Decoding of Auditory Attention to Conversations with Turn-taking Speakers"

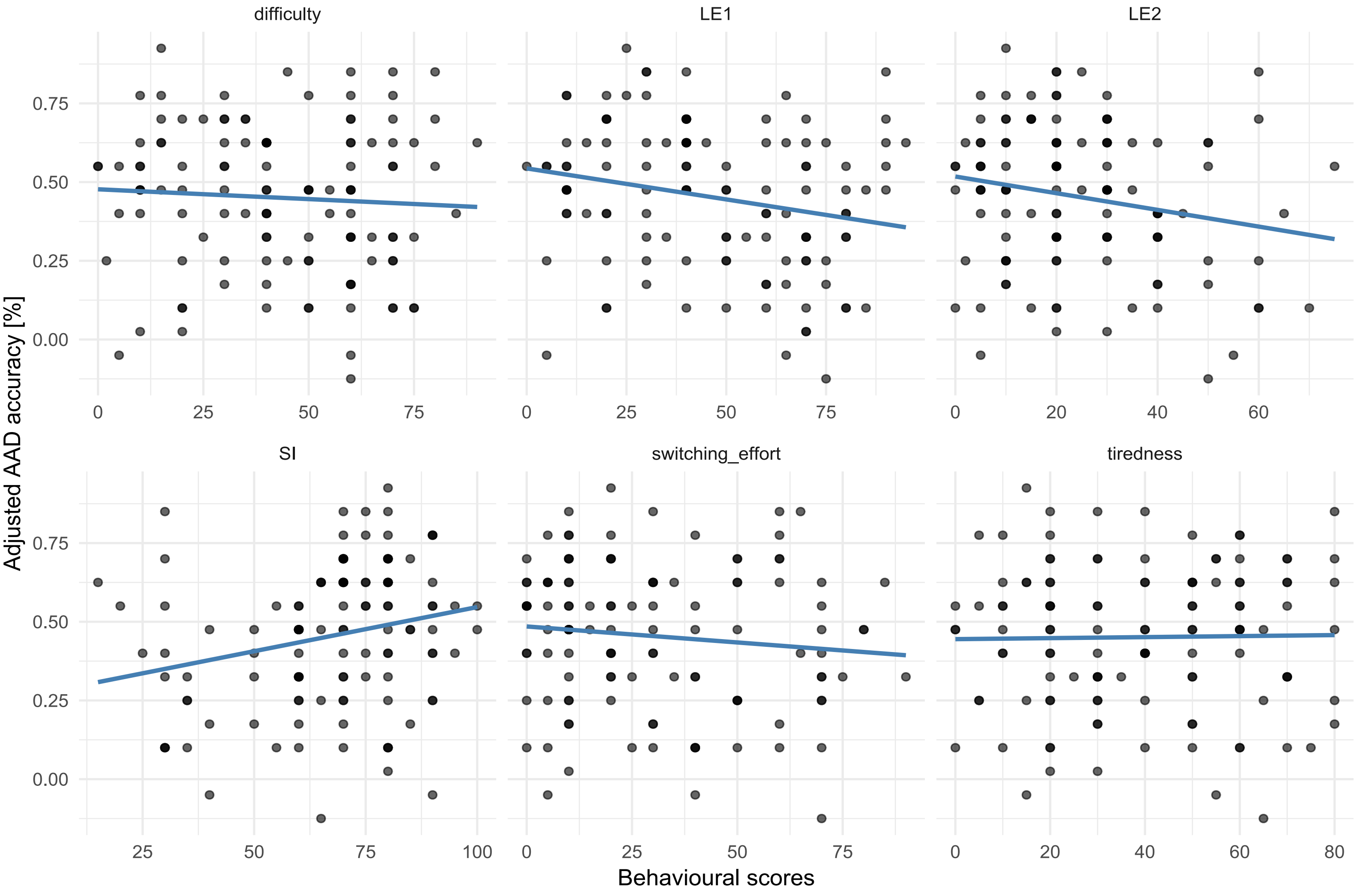


*Supplementary Figure 1: Scatterplots of the different behavioural measures. Behavioural scores are shown in function of adjusted accuracy*
